# Network-based representation learning enables the identification of risk genes and latent functional pathways in primary open-angle glaucoma

**DOI:** 10.1101/2024.08.15.608134

**Authors:** Henry C. Cousins, Russ B. Altman, Louis R. Pasquale

## Abstract

Despite the identification of hundreds of risk genes for primary open-angle glaucoma (POAG), a significant portion of the POAG genetic risk landscape remains unexplored. We hypothesized that unsupervised learning on large protein-protein interaction (PPI) networks could enable comprehensive characterization of the genetic pathways that underlie POAG risk. We used graph representation learning on a proteome-scale PPI network to generate embeddings capturing complex features of each protein’s interactions. Using these embeddings, we trained a model with POAG-associated genes from the DisGeNET database to output an inferred POAG risk score for over 12,000 gene products, which identified known POAG risk genes with an area under the receiver operating characteristic curve of 0.739 (95% CI 0.686-0.792). These included well-known POAG risk genes such as *RHOA* and *MMP3*, as well as genes with significant contributions to other ocular diseases. Pathway analysis on the proteome-wide risk scores implicated 20 biological processes in POAG pathogenesis. Furthermore, cluster analysis of embeddings for POAG risk genes revealed 5 distinct functional neighborhoods, including cytokine signaling, coagulation response, collagen biosynthesis, extracellular matrix development, and fatty acid metabolism. Our results suggest that representation learning can recognize important patterns of protein interaction that allow *in silico* prioritization of POAG risk genes and pathways.

## INTRODUCTION

Primary open-angle glaucoma (POAG) is the world’s leading cause of permanent blindness.^1^ With an estimated global prevalence of 3.5% and an age-dependent pattern of incidence, the global disease burden of POAG will expand significantly in the coming decades, with projections exceeding 100 million affected individuals by 2040.^2^ The insidious nature of the disease’s morbidity arises from its progressive, irreversible course, whereby heterogeneous risk factors promote chronic damage to the optic nerve.^3^ While environmental factors represent key risk factors for the disease, much of the pathogenesis of POAG depends on genotype, with heritability estimates approaching 70%.^4,5^

Nonetheless, the specific patterns of genomic variation that underlie POAG risk remain incompletely understood. Genetic linkage studies originally identified a small number of genes with high effect on POAG risk, including *OPTN, MYOC*, and *TBK1*.^6–9^ These genes were principally identified in small genetic linkage studies with fewer than 500 patients. However, in the past decade, large genome-wide associations studies (GWAS), themselves enabled by the construction of population-scale biobanks, have enabled mapping of the genetic basis of POAG in far greater detail.^10–17^ To date, GWAS analyses have identified over 300 distinct risk genes for POAG, although at least 90% of known disease heritability likely remains unexplained.^11^

Known POAG risk genes to date fall into several distinct functional categories. Genes involved in extracellular matrix biosynthesis are particularly well represented in GWAS analyses of both POAG itself and related endophenotypes such as intraocular pressure (IOP) and cup-to-disc ratio.^12,18^ Such genes, including those encoding various collagen chains (*COL4A3, COL5A1, COL8A2*, and *COL8A1*) and metalloproteinases (*ADAMTS8* and *ADAMTS2*), may affect IOP directly by modulating trabecular meshwork structure and outflow resistance.^19–22^ Several genes involved in regulation of vascular tone and lipid metabolism, including *NOS3, ABCA1, ARHGEF12*, and Rho-family GTPases, also have strong associations with POAG, which may relate to the importance of these pathways in IOP regulation.^23,24^ *CAV1* and *CAV2*, among the most prominent POAG risk genes, directly regulate microvascular tone by modulating nitric oxide signaling.^17,23,25^ Finally, a large number of POAG-associated genes function broadly as regulators of neuroinflammation, including *IL2, CXCL5*, and *TNF-ɑ*, as well as *TBK1* and *OPTN*, mutations in which are causal for certain glaucoma presentations.^23,26,27^

Recent improvements in resolving the genome-wide POAG risk landscape have enabled profound clinical advances. Polygenic risk scores, previously considered impractical in glaucoma, have begun to approach clinical viability as prognostic tools.^28,29^ Furthermore, genomic insights have directly enabled an emerging paradigm of rational drug development in glaucoma, exemplified by the discovery of Rho kinase inhibitors and adenosine receptor agonists as protective agents in POAG.^30,31^ Nonetheless, the genome-wide landscape of POAG risk remains poorly understood, with most heritability unaccounted for by known risk genes.^11^

The development of large-scale protein-protein interaction (PPI) databases and efficient machine-learning techniques for distilling complex genetic data offers a means of identifying underlying genomic signatures of POAG pathogenesis. Specifically, unsupervised learning on large PPI networks enables individual genes to be represented in a common embedding space reflecting complex interactions among gene products.^32,33^ Because genes with similar biochemical functions have similar embeddings under this paradigm, functional insights from limited experimental data can provide a basis for more complete annotation of the genome with respect to disease risk.^34^ This bioinformatic premise has demonstrated an ability to resolve new pathogenic genes in cancer, coronary artery disease, and Alzheimer’s disease, among other conditions.^35–39^

We hypothesized that POAG risk genes may participate in complex biochemical processes that are not obvious from individual interactions alone. These processes allude to an underlying genetic architecture of the disease based on complex interactions among many gene products. To infer this latent genetic architecture, we developed a model based on low-dimensional PPI embeddings of POAG-associated gene products (**Figure 1**). We demonstrate that this approach enables both prediction of disease risk for individual genes and inference of genetic pathways that may contribute to POAG pathogenesis.

**Figure 1.**
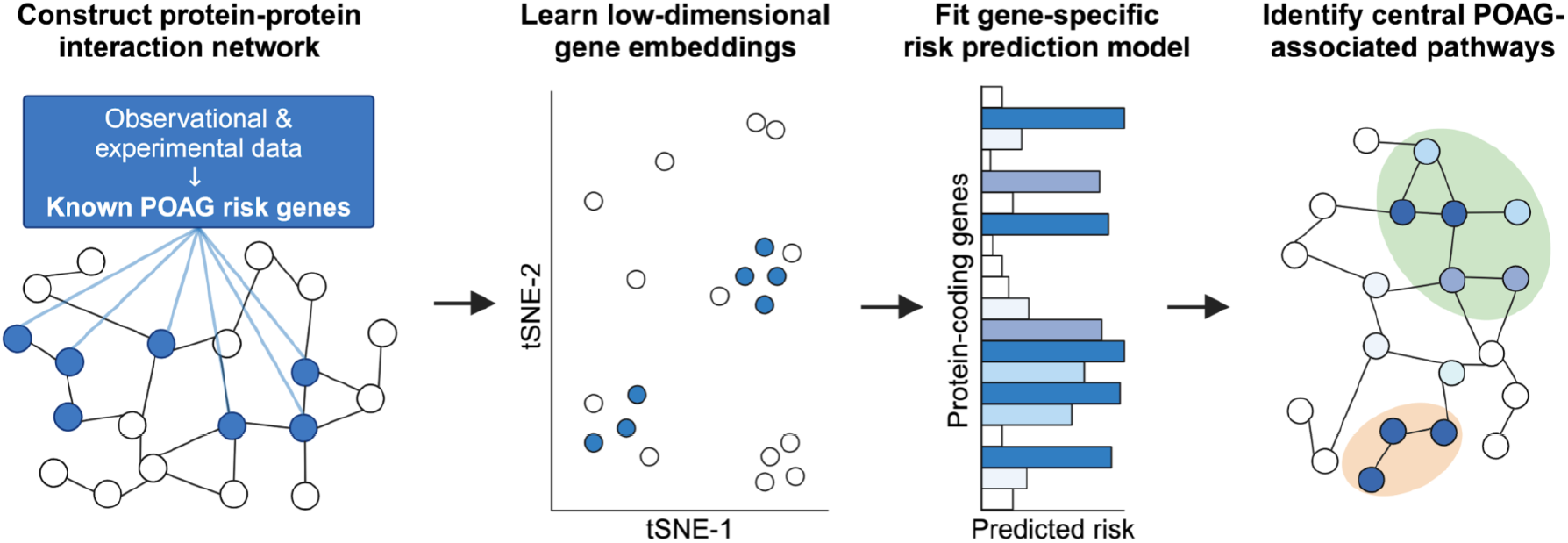
Overview of POAG risk gene embedding approach. A proteome-scale PPI network was constructed using high-confidence protein-protein interactions. Known gene associations with POAG were obtained from GWAS and experimental sources. Next, unsupervised representation learning enables the calculation of low-dimensional node embeddings that represent each gene product’s local and global context in the PPI network. We then fit a supervised model using the known POAG associations to learn a risk score for each gene, from which a variety of downstream analyses, including the resolution of central pathways, are possible.

## METHODS

### Generating embeddings

To generate the PPI network, we first obtained high-confidence protein interactions from the STRING database (version 11).^33^ We used these interactions to construct an undirected network representing all high-confidence interactions among proteins in the human proteome. Proteins lacking known high-confidence interactions with other proteins were excluded. The complete network contained 12,396 distinct proteins (“nodes”) and 324,152 interactions (“edges”) among them.

We next generated low-dimensional embeddings for each node in the full network using the node2vec algorithm.^40^ Node2vec is a two-part algorithm that first generates random walks through edges in the network, using a weighting bias to preserve local architecture, followed by a neural-network-based encoding of the nodes within each walk. For each protein in the full PPI network, we obtained in this way a 128-length vector embedding that reflects its interactions with other proteins, with similar embeddings connoting similar functional relationships. To visualize the node2vec embeddings, we next performed t-distributed stochastic neighbor embedding (tSNE) from the original 128 dimensions into a 2-dimensional space.

### Training model

We next obtained the set of genes with known POAG association in the DisGeNET database, which includes evidence from both GWAS and experimental sources.^41^ 383 genes with a high-confidence POAG association were identified, of which 294 encoded gene products with representation in the PPI network.

We next fit a supervised learning model on the node2vec embeddings to model the relationship between PPI context and POAG risk, mapping each raw embedding to a probability of POAG association. We used an L2-regularized logistic regression (LR) model, trained using Monte Carlo cross-validation for 100 iterations, with 80% of the data used for training and 20% for evaluation. To calculate individual risk predictions for each gene, we created an ensemble model based on the average of all predictions for each gene. We measured the performance of the model using the area under the receiver operating characteristic (AUROC), as well as a confusion matrix.

### Pathway analysis

To assess the representation of known functional pathways in the model’s proteome-wide predictions, we next performed the prerank variation of gene set enrichment analysis on the complete list of gene-wise predictions, using gene ontology (GO) biological process (BP) gene sets (2023 release).^42–44^ We considered all gene sets with at least 5 members, interpreting significance as a family-wise error rate (FWER) less than 0.05.

### Embedding cluster analysis

To determine whether the POAG gene embedding space could reveal an underlying structure to the risk gene landscape, we next performed a cluster analysis to identify latent groupings of risk genes. We first used the elbow method to identify an optimal set of clusters within the set of embeddings for all known POAG risk genes. We next performed k-means clustering on the full set of risk gene embeddings to define distinct clusters of gene embeddings. Finally, to assign semantic interpretation to the distinct embedding clusters, we performed an overrepresentation analysis on each cluster using GO-BP terms.

### Computational resources

All analyses were performed in Python 3.10 using the NumPy 1.25.2, pandas 2.0.3, scikit-learn 1.2.2, SciPy 1.11.4, seaborn 0.13.1, and gseapy 1.1.3 packages in addition to the standard library.

## RESULTS

We first generated protein-specific embeddings from the complete PPI network using node2vec, intended to capture local and global interaction data for each of the 12,396 proteins in the network. The resulting embeddings showed a significant tendency to cluster, with a Hopkins statistic of 0.89, consistent with their organization into discrete functional neighborhoods (**Figure 2**).

**Figure 2.**
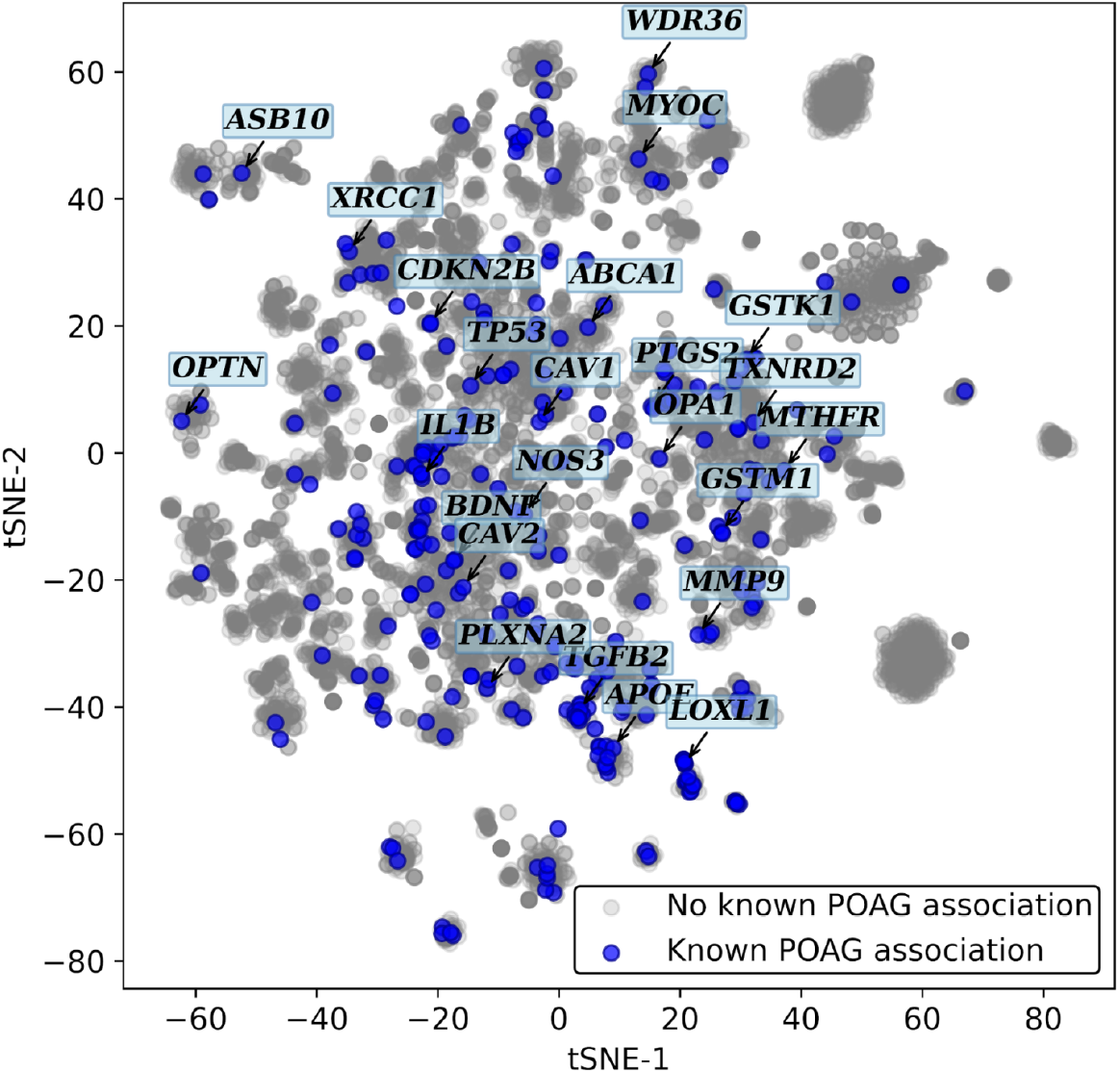
POAG risk genes colocalize in PPI-derived embedding space. Visualization of full PPI network embeddings using tSNE projections, showing distinct clustering among gene products. POAG-associated genes (blue) tend to colocalize in the embedding space, suggesting that disease genes occupy distinct functional neighborhoods.

In parallel, we identified 294 genes with a known POAG association in DisGeNET. This collection included many well-known POAG risk genes, including *OPTN, TBK1, CAV1*, and *ADAMTS10*. Of note, known POAG risk genes tended to have high centrality in the PPI network compared to other genes, with a mean normalized degree centrality of 0.011 (SD 0.014) for POAG risk genes versus 0.0084 (SD 0.013) for other genes (p < 0.001).

The trained logistic regression model identified known POAG-associated risk genes with a mean AUROC of 0.739 (95% CI 0.686-0.792) (**Figure 3A**). The highest-scoring genes included several with well-known associations with POAG, including *TP53, PTGS2, RHOA, VEGFA*, and *MMP3*. High-scoring genes without known POAG associations included *HSP90AA1, PTGES3, AR, STAT6*, and *IL13* (**Figure 3B, Table I**).

**Table I.** Individual genes with highest predicted risk scores.

| Rank | Name | Function | Model score | Known POAG association |
| --- | --- | --- | --- | --- |
| 1 | *TP53* | Tumor suppressor | 71.1 | Yes |
| 2 | *HSP90AA1* | Protein folding | 67.5 | No |
| 3 | *PTGES3* | Prostaglandin synthesis | 56.7 | No |
| 4 | *AR* | Androgen signaling | 50.1 | No |
| 5 | *STAT6* | Cytokine signaling | 48.6 | No |
| 6 | *PTGS2* | Prostaglandin synthesis | 46.9 | Yes |
| 7 | *IL13* | Cytokine signaling | 46.3 | No |
| 8 | *ALOX5* | Leukotriene synthesis | 46.0 | No |
| 9 | *RHOA* | Signal transduction | 44.7 | Yes |
| 10 | *VEGFA* | Angiogenesis | 43.7 | Yes |

**Figure 3.**
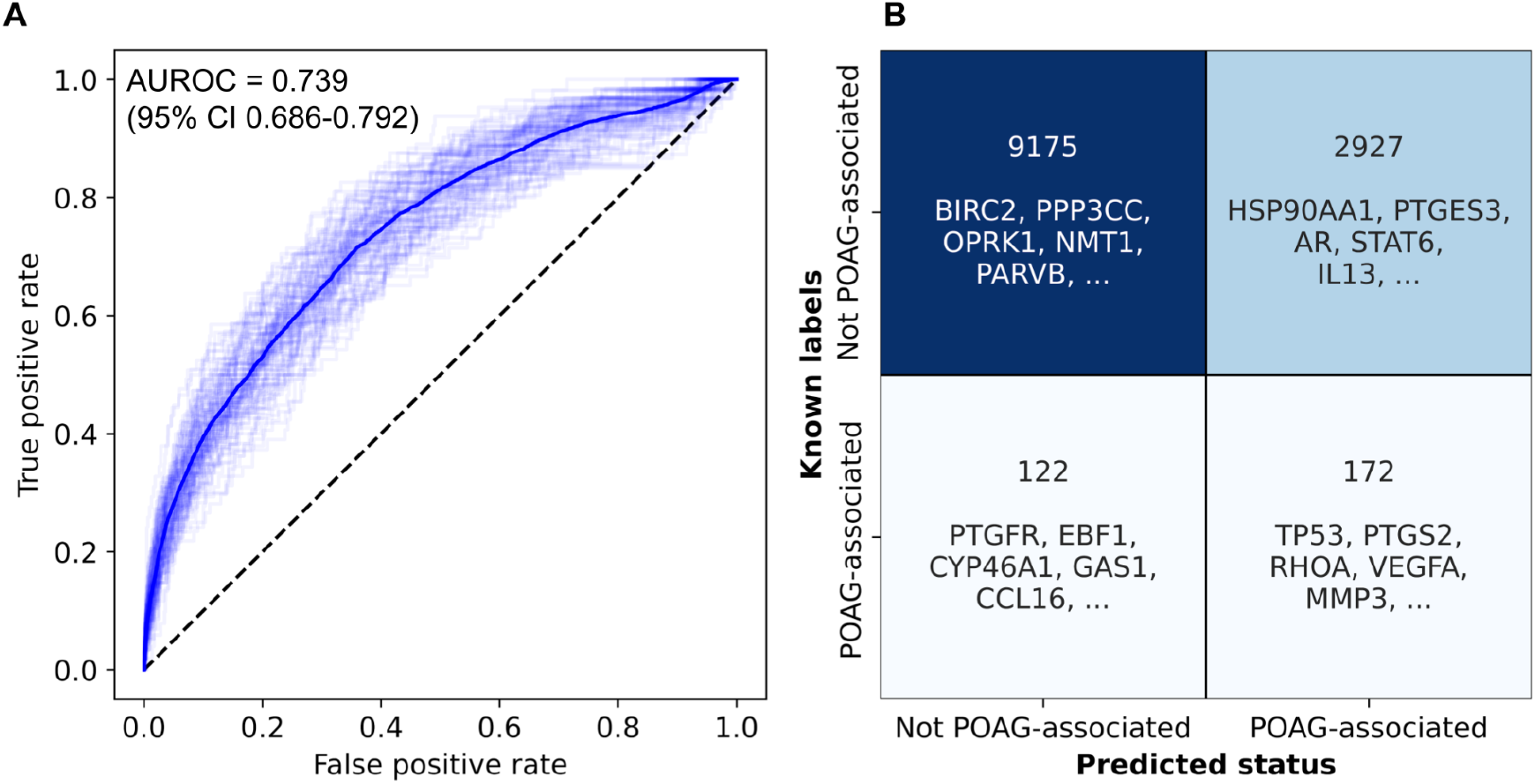
Supervised learning on PPI embeddings enables retrieval of known POAG-associated genes. (A) ROC curve for identification of known POAG-associated genes. The AUROC of the model was 0.739 (95% CI 0.686-0.792). (B) Confusion matrix for model predictions, compared to literature-derived labels. True positives include well-studied POAG-associated genes including *RHOA, VEGFA*, and *MMP3*. Of note, predicted positives without a known POAG association, including the prostaglandin synthase *PTGES3* and the transcription factor *STAT6*, tended to have associations with ocular inflammatory processes, suggesting a molecular basis for involvement in POAG.

Next, we measured the enrichment of GO-BP gene sets in the proteome-wide list of gene scores. 20 gene sets were positively enriched at an adjusted p-value threshold of 0.05 (**Figure 4**). These included multiple pathways related to extracellular matrix and collagen biosynthesis (GO:0045229, GO:0043062, GO:0030198, GO:0030199, all p < 0.001), arachidonic acid and icosanoid metabolism (GO:0006690 and GO:0019369, p < 0.001), endothelial cell maturation (GO:0035987, p = 0.004; GO:2000352, p = 0.009; GO:0043536, p = 0.04) and coagulation pathways (GO:0030195, p = 0.003; GO:1904407, p = 0.004; GO:0045429, p = 0.004).

**Figure 4.**
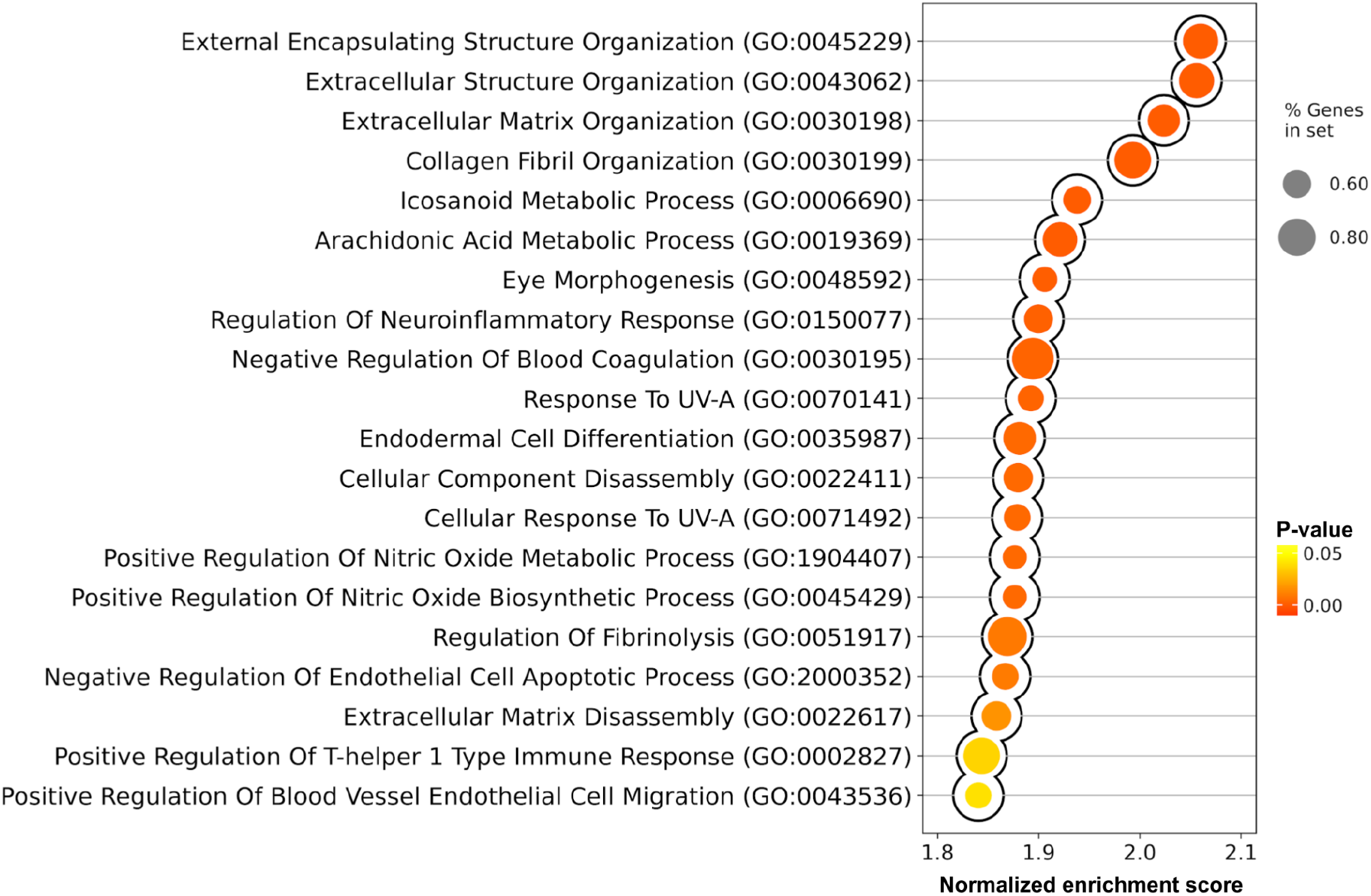
Top enriched pathways among predicted proteome-wide gene risks. Twenty gene sets were significantly enriched by gene set enrichment analysis. Gene set enrichment analysis of the proteome-wide risk predictions reveal enrichment of extracellular matrix biosynthesis pathways. Fatty acid metabolism and nitric oxide signaling were also enriched, consistent with their importance in early disease pathogenesis.

Finally, we asked whether the geometry of the embeddings for POAG risk genes could reveal underlying relationships among risk genes that would not otherwise be evident. Specifically, we performed cluster analysis of the embeddings for all predicted POAG risk genes, using k-means clustering with k of 5, determined using the elbow method. To characterize the function of each of the five clusters, we used overrepresentation analysis to identify the most well-represented GO-BP terms in each cluster (**Figure 5A**). Each cluster was largely distinct in terms of its constituent pathways, indicating uniqueness in the respective biological functions of each cluster (**Figure 5B**). Cluster 1 corresponded to genes involved with cytokine signaling and leukocyte differentiation, Cluster 2 to the coagulation response, Clusters 3 and 4 to collagen and extracellular matrix development, and Cluster 5 to fatty acid metabolism.

**Figure 5.**
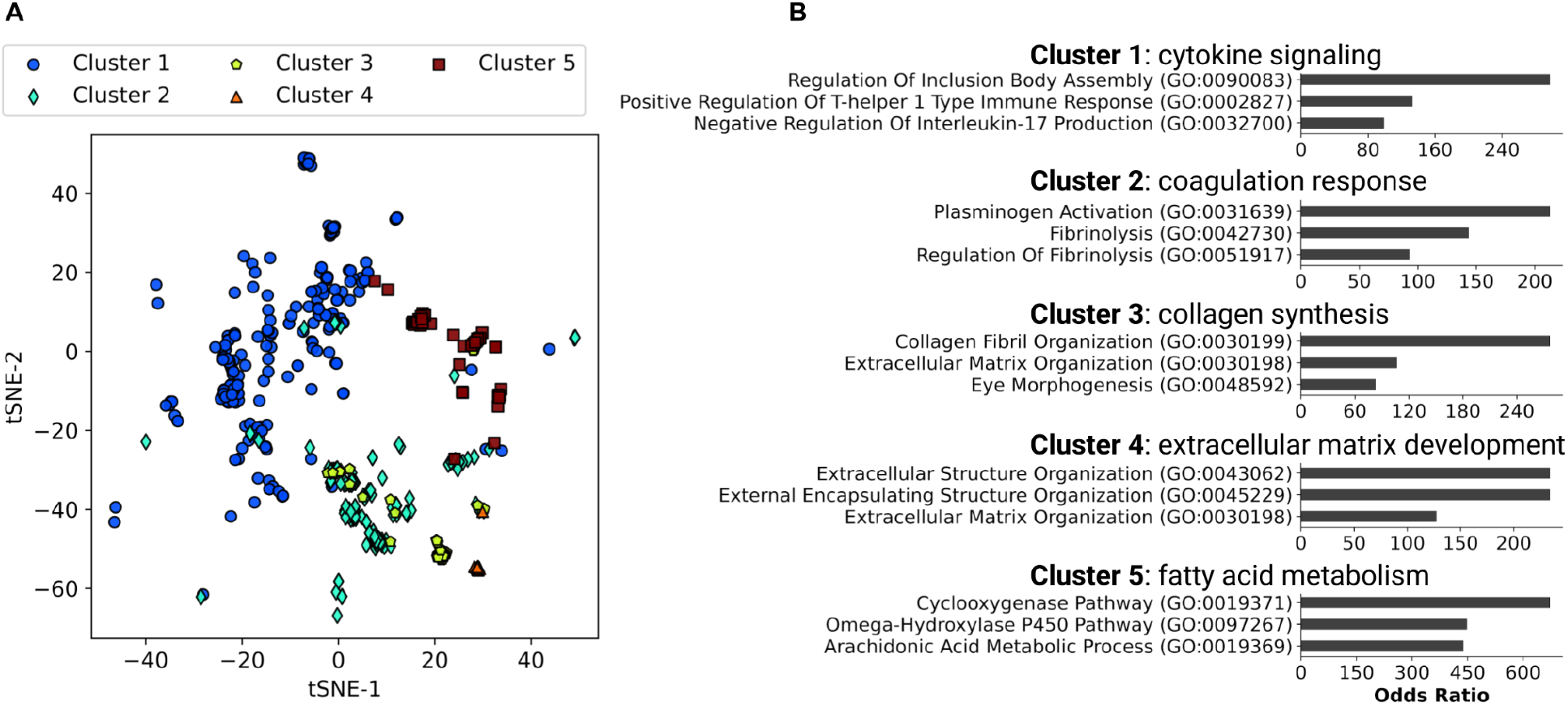
Embedding cluster analysis of predicted POAG risk genes. (A) Predicted POAG-associated genes comprised 5 distinct clusters, as identified by k-means. (B) GO-BP enrichment analysis revealed distinct biological functions for each cluster (top 3 most significant terms shown for each cluster). Cluster 1 corresponded to genes involved with cytokine signaling and leukocyte differentiation, Cluster 2 to the coagulation response, Clusters 3 and 4 to collagen and extracellular matrix development, and Cluster 5 to fatty acid metabolism.

## CONCLUSIONS

Our work demonstrates that PPI networks contain sufficient information to infer POAG associations for genes whose empiric disease associations remain to be characterized fully. Importantly, such predictive power would not be possible without network-based unsupervised learning methods and large PPI collections. The model implicates both well-studied POAG risk factors such as prostaglandin and Rho GTPase signaling as well as genes involved in other ocular pathologies, such as VEGF and IL13 pathways.^23,45^ By characterizing the risk landscape throughout the human proteome, we perform a more comprehensive analysis of POAG risk pathways than has previously been possible. Furthermore, we show that POAG risk genes are not randomly distributed but tend to cluster into functional neighborhoods, which are well visualized in high-dimensional embedding space representing PPI relationships. These neighborhoods imply a core role for genes related to cytokine signaling, coagulation, collagen synthesis, ECM development, and fatty acid metabolism, which may help to focus subsequent investigations of individual gene function.

In addition to recapitulating known POAG risk genes, we also identified several genes that scored highly in our model but lacked a known empiric association with POAG. The highest-scoring such gene was *HSP90AA1*, encoding a widely expressed heat-shock protein induced in response to proteotoxic stress. Intriguingly, HSP90AA1 directly modulates inflammatory signaling via TGF-alpha and NF-kB, both of which play a critical role in ocular hypertension.^46–48^ Although most well documented in breast cancer, *HSP90AA1* polymorphisms are also associated with neurodegenerative conditions like Alzheimer’s disease and Parkinson’s disease, likely via dysregulated protein chaperoning.^49^ Another highly scoring gene without a known POAG association was *PTGES3*, which facilitates synthesis of prostaglandin E2, which itself directly increases aqueous humor outflow.^50^ The model also implicated *STAT6*, which facilitates the IL4-mediated inflammatory response and has been associated with chronic ocular inflammation in keratoconjunctivitis and blepharitis.^51^

More broadly, our work demonstrates a new approach toward risk gene prioritization in glaucoma. Although recent large GWAS have vastly broadened the known landscape of POAG risk genes, they remain limited in their ability to identify risk genes with infrequent variation.^12^ While differential gene expression analysis can supplement these deficiencies to some extent, they are sensitive to experimental noise and challenging to perform on large patient cohorts.^52^ Purely bioinformatic approaches, such as ours, provide a useful addition due to their scale, efficiency, and interpretability. Specifically, by applying unsupervised learning to large protein interaction databases, we generate concise, expressive representations of individual genes that can augment limited experimental data to reveal more disease-relevant patterns.

A principal limitation of PPI-based methods such as ours lies in their reliance on literature-derived PPI collections, which may emphasize well-studied proteins at the expense of others. Furthermore, PPI collections like STRING do not delineate between common, functional interactions between proteins and those that may be rarer in nature or less biologically relevant. Future efforts to improve on our approach may ameliorate some of these issues through the use of new cell- or disease-specific PPI collections.

In conclusion, we demonstrate that unsupervised learning on protein interaction networks enables proteome-scale inference of risk genes in open-angle glaucoma. In addition to providing systematic inference of POAG associations for both individual genes and biological pathways, this approach also enables resolution of underlying functional clusters of risk genes. As a whole, our results suggest that the combination of representation learning and large protein interaction datasets may augment existing experimental methods in explaining the complex genetic architecture of POAG.

